# CLN3 regulates endosomal function by modulating Rab7A activity

**DOI:** 10.1101/634915

**Authors:** Seda Yasa, Graziana Modica, Etienne Sauvageau, Abuzar Kaleem, Guido Hermey, Stephane Lefrancois

## Abstract

Mutations in *CLN3* are a cause of juvenile NCL (JNCL), also known as Batten Disease. Clinical manifestations includes cognitive regression, progressive loss of vision and motor function, epileptic seizures, and a significantly reduced lifespan. CLN3 localizes to endosomes and lysosomes, and has been implicated in intracellular trafficking and autophagy. However, the precise molecular function of CLN3 remains to be elucidated. We show that CLN3 interacts with Rab7A, a small GTPase that regulates several functions at late endosomes. We found that CLN3 is required for the efficient endosome-to-TGN trafficking of the lysosomal sorting receptors by regulating the Rab7A/retromer interaction. In cells lacking CLN3 or expressing CLN3 harbouring a disease-causing mutation, the lysosomal sorting receptors were degraded. We also demonstrated that CLN3 is required for the Rab7A/PLEKHM1 interaction, which is required for autophagosome/lysosome fusion. Overall, our data provides a molecular explanation behind phenotypes observed in JNCL.

## Introduction

The Neuronal Ceroid Lipofuscinoses (NCLs) are a group of rare neurodegenerative diseases linked to over 430 mutations in 13 genetically distinct genes (*CLN1-8, CLN10-14*) (Mole and Cotman, 2015). Clinical manifestations of NCLs include intellectual impairment, progressive loss of vision and motor function, epileptic seizures, and a significantly reduced lifespan (Anderson et al., 2012). At the cellular level, NCLs display aberrant lysosomal function and an excessive accumulation of ceroid lipofuscin in neurons as well as other cell types outside of the central nervous system (Anderson et al., 2012).

Juvenile Neuronal Ceroid Lipofuscinosis (JNCL) is caused by germline mutations in Ceroid lipofuscinosis neuronal-3 (CLN3). More commonly referred to as Batten disease, it is the most common paediatric neurodegenerative disease (Anderson et al., 2012; Mole and Cotman, 2015). CLN3 is a protein of 438 amino acids with six transmembrane domains whose N- and C-terminal ends are located in the cytosol (Ratajczak et al., 2014). CLN3 is a highly glycosylated integral membrane protein (Storch et al., 2004) that localizes to the endosomal/lysosomal membrane (Oetjen et al., 2016) among other intracellular locations and has proposed roles in lysosomal trafficking and autophagy (Lojewski et al., 2014; Metcalf et al., 2008). CLN3 interacts with and is implicated in Rab7A recruitment to endosomal membranes (Uusi-Rauva et al., 2008), however the function of this interaction is unknown. Furthermore, how disease-causing mutations in CLN3 affect this interaction or downstream functions of Rab7A has not been elucidated.

The <u>Ra</u>s-like proteins in <u>b</u>rain (Rabs) are key regulators of the formation, trafficking, and fusion of transport vesicles at the endoplasmic reticulum (ER), Golgi apparatus and early and late endosomes (Hutagalung and Novick, 2011). Rabs function by interacting with downstream effectors (Grosshans et al., 2006) and a key process regulating these interactions is the GTP loading or “activation” of Rab GTPases at specific membrane sites (Pfeffer and Aivazian, 2004). This GDP to GTP switch is regulated by guanine exchange factors (GEFs) that load Rab GTPases with GTP, while GTPase activating proteins (GAPs) terminate their activity by hydrolyzing the GTP to GDP (Barr and Lambright, 2010).

Active Rab7A localizes to endosomal membranes and recruits numerous effectors to perform a variety of different functions such as endosome-to-trans Golgi Network (TGN) trafficking (Rojas et al., 2008; Seaman et al., 2009), autophagosome/lysosome fusion (McEwan et al., 2015), lysosomal positioning (Wijdeven et al., 2016) and degradation of endocytic cargo such as epidermal growth factor (EGF) receptor (EGFR) (Vanlandingham and Ceresa, 2009). In this study, we systematically analyzed several of these Rab7A mediated pathways to determine which was under the control of CLN3. We identified defects in endosome-to-TGN sorting, EGFR degradation and autophagy. Overall, our data suggests a role for CLN3 as an upstream regulator of Rab7A function.

## Results

### Disease-causing mutations in CLN3 increases its interaction with Rab7A

It has been reported that CLN3 interacts with the small GTPase Rab7A (Uusi-Rauva et al., 2012), but the functional role of this interaction is not understood nor how disease-causing mutations in CLN3 affect this interaction. We used bioluminescence resonance energy transfer (BRET) to confirm the CLN3/Rab7A interaction in live cells and to determine how disease-causing mutations affect this interaction. Compared to co-immunoprecipitation, BRET is performed in live cells, with proteins localized to their native environment. From BRET titration curves, the BRET_50_ can be calculated which is the value at which the concentration of the acceptor is required to obtain 50% of the maximal BRET signal (BRET_MAX_) and is indicative of the propensity to interact (Kobayashi et al., 2009; Mercier et al., 2002). The smaller the BRET_50_ the stronger the interaction. Renilla Luciferase II (RlucII) was fused at the N-terminus to wild-type Rab7A (RlucII-Rab7A). As previously shown, this tag had little effect on the distribution or function of Rab7A as expressing RlucII-Rab7A in Rab7A knockout cells (Rab7A-KO) rescued retromer recruitment (Modica et al., 2017). The energy acceptor green fluorescence protein 10 (GFP10) was fused at the N-terminus of wild-type and various point mutation containing CLN3 (GFP10-CLN3, GFP10-CLN3^R334H^, GFP10-CLN3^V330F^, GFP10-CLN3^E295K^, GFP10-CLN3^L101^P). We engineered a CLN3 mutant harbouring the common exon 7 and 8 deletion (CLN3^Δex7-8^), but this protein was not efficiently expressed in our hands. HeLa cells were co-transfected with a constant amount of RlucII-Rab7A and increasing amounts of GFP10-CLN3 to generate BRET titration curves. The BRET signal between RlucII-Rab7A and GFP10-CLN3 rapidly increased with increasing amounts of expressed GFP10-CLN3 until it reached saturation, suggesting a specific interaction (**Figure 1a, blue curve**). We also tested another Rab GTPase, Rab1a, which is localized to the Golgi (Dumaresq-Doiron et al., 2010) to confirm the specificity of the CLN3/Rab7A interaction. We generated BRET titration curves with RlucII-Rab1a (**Figure 1a, red curve**) in HeLa cells. We extrapolated the BRET_50_ for the interaction between CLN3 and the two Rab GTPases and found that the Rab7A/CLN3 interaction had a much smaller BRET_50_ (0.004) compared to the BRET_50_ (0.020) Rab1a/CLN3 interaction, indicating Rab7A has a higher propensity to interact with CLN3 compared to Rab1a (**Figure 1b**). A small fraction of CLN3 localizes to the Golgi (Cao et al., 2006), hence we are not surprised that we detected an interaction between CLN3 and Rab1a. We next performed a BRET experiment to determine the impact of CLN3 disease-causing mutations on the CLN3/Rab7A interaction (**Figure 1c**). Compared to wild-type CLN3, we found a significantly stronger (smaller BRET_50_) interaction between Rab7A and CLN3^R334H^ and CLN3^V330F^ (**Figure 1d**), while the CLN3^E295K^ and CLN3^L101P^ mutations had negligible affects on the CLN3/Rab7A interaction (**Figure 1d**).

**Figure 1.**
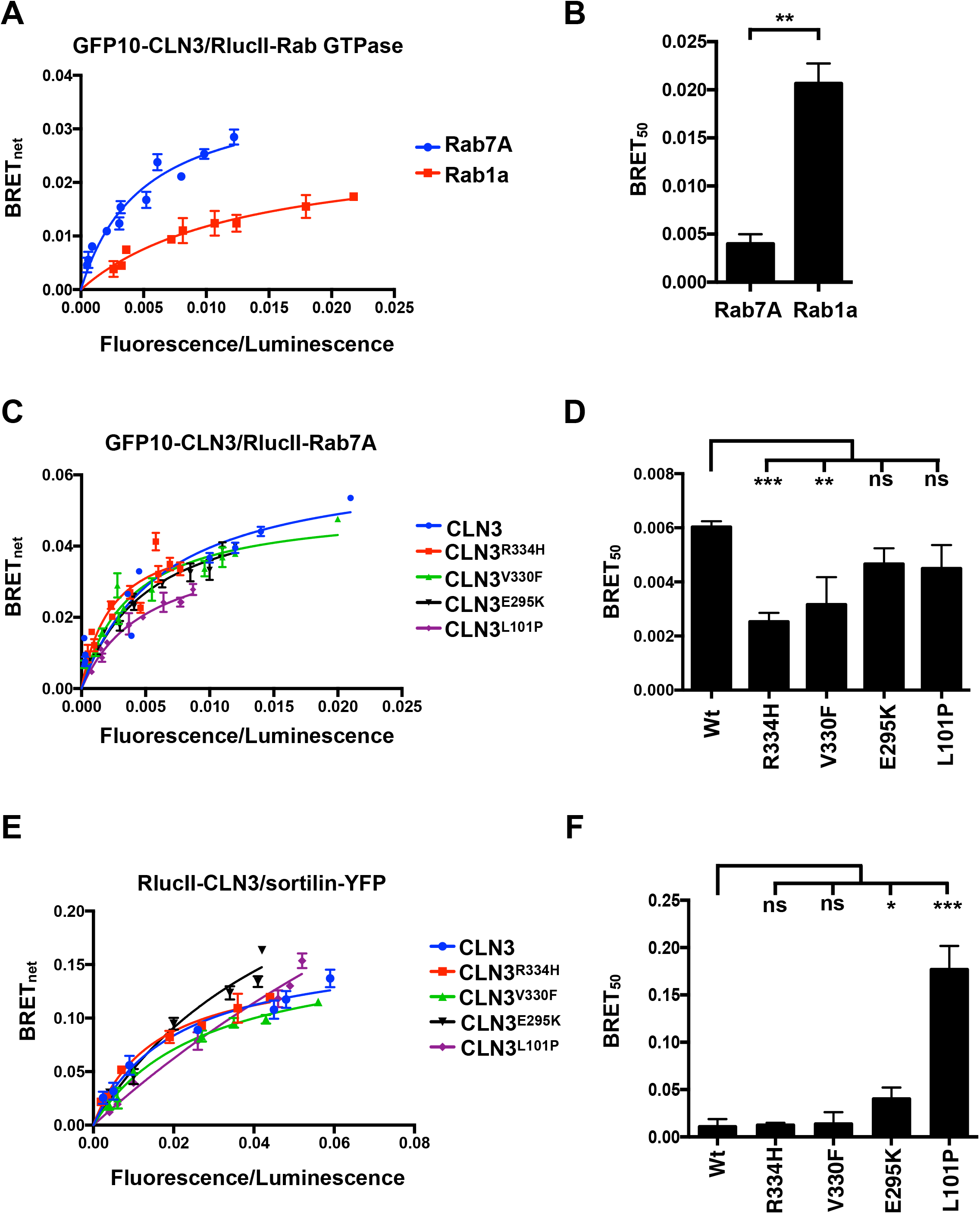
CLN3 interacts with Rab7A and sortilin. (A) HeLa cells were transfected with a constant amount of RlucII-Rab7A or RlucII-Rab1a and increasing amounts of GFP10-CLN3 to generate BRET titration curves. 48 hours posttransfection, BRET analysis was performed. BRET signals are plotted as a function of the ratio between the GFP10 fluorescence over RlucII luminescence. (B) BRET_50_ was extrapolated from three independent experiments. Data is represented as mean ± SD. **, P ≤ 0.01; Student’s t-test. (C) Hela cells were transfected with a constant amount of RlucII-Rab7A and increasing amounts of GFP10-CLN3, GFP10-CLN3^R334H^, GFP10-CLN3^V330F^, GFP10-CLN3^E295K^, or GFP10-CLN3^L101P^ to generate BRET titration curves. BRET_MAX_: maximal BRET signal. (D) BRET_50_ was extrapolated from 3 three independent experiments. Data is represented as mean ± SD. ns, not significant; **, P ≤ 0.01; ***, P ≤ 0.001; two-way ANOVA followed by Tukey’s post hoc test. (E) HeLa cells were transfected with a constant amount of RlucII-CLN3 or RlucII-CLN3 harbouring a disease-causing mutation and increasing amounts of sortilin-YFP to generate BRET titration curves. 48 hours post-transfection, BRET analysis was performed. BRET signals are plotted as a function of the ratio between the YFP fluorescence over RlucII luminescence. (F) BRET_50_ was extrapolated from 3 three independent experiments. Data is represented as mean ± SD. ns, not significant; **, P ≤ 0.01; ***, P ≤ 0.001; two-way ANOVA followed by Tukey’s post hoc test.

Mutations in CLN5, which interacts with CLN3, cause a form of NCL with overlapping symptoms to CLN3 disease (Vesa et al., 2002). In our previous work, we showed that CLN5 interacts with sortilin (Mamo et al., 2012). Therefore, we rationalized that CLN3 could also interact with sortilin. To test this hypothesis, we performed a BRET experiment by expressing a constant amount of RlucII-tagged wild-type CLN3 (RlucII-CLN3) and increasing amounts of yellow fluorescence protein (YFP) tagged sortilin (sortilin-YFP). We found an interaction between CLN3 and sortilin, as the curve reached saturation (**Figure 1e, blue curve**). We next determined the impact CLN3 disease-causing mutations could have on this interaction. We generated BRET titration curves between sortilin-YFP and RlucII-CLN3^R334H^ (**Figure 1e, red curve**), RlucII-CLN3^V330F^ (**Figure 1e, green curve**), RlucII-CLN3^E295K^ (**Figure 1e, black curve**) and RlucII-CLN3^L101P^ (**Figure 1e, purple curve**). Compared to wild-type CLN3, we found a significant increase in the BRET_50_ values for the sortilin/CLN3^E295K^ and sortilin/CLN3^L101P^ interactions, suggesting weaker interactions (**Figure 1f**). Interestingly, we found no change in the sortilin/CLN3^R334H^ and sortilin/CLN3^V330F^ interactions compared to the sortilin/CLN3 interaction (**Figure 1f**).

A previous study had found that Rab7A immunofluorescence staining was decreased in cells harbouring the homozygous mutation (CLN3^Δex7-8^/CLN3^Δex7-8^) compared to wild-type cells (Fossale et al., 2004). Based on this, we investigated whether CLN3 is required for the membrane recruitment of Rab7A by generating CLN3-KO HeLa cells using CRISPR/Cas9 (**Figure S1a**). We also generated Rab7A-KO cells using the same parental HeLa strain to serve as a control in our experiments (**Figure S1b**). Previous work has demonstrated that cells harbouring CLN3 mutations accumulate LC3II-positive autophagosomes (Fossale et al., 2004; Vidal-Donet et al., 2013). Further studies demonstrated that CLN3 mutations affected downstream Rab7A functions, but the molecular details were not determined (Chandrachud et al., 2015). Soluble microtubule-associated protein 1A/1B-light chain 3 (LC3) forms LC3-II through lipidation upon activation of autophagy (Kominami et al., 2008). LC3-II then localizes to autophagosomal membranes and remains there until the autophagosomes fuse with the lysosomes, resulting in its degradation (Ganley et al., 2011). If autophagosome/lysosome fusion is inhibited, or if lysosomal function is blocked either by a compound such a bafilomycin A1 (BafA1) or a defective protein, degradation of LC3-II can not occur. Therefore, comparing the turnover of LC3-II levels is a good measure of autophagic activity and competence. To test if our CLN3-KO cells behaved like as expected with respect to autophagy, we initiated autophagy through nutrient starvation by incubating wild-type, CLN3-KO and Rab7A-KO cells with EBSS media in the presence or not of BafA1, a compound known to inhibit lysosomal function. We found an increase in LC3-II/Total LC3 levels in wild-type cells upon starvation compared to fed conditions, which was further increased in BafA1 treated cells (**Figure S1c and d**) suggesting wild-type HeLa cells could initiate LC3 recruitment and degrade it, unless lysosomal function was inhibited pharmacologically. Next, we used Rab7A-KO HeLa cells as a control group, since without Rab7A, we would expect defects in LC3 degradation and therefore higher a ratio of LC3-II/total LC3 (**Figure S1c and d**). As expected, we found higher basal levels of LC3-II/Total LC3 levels, which increased upon the induction of autophagy. The addition of BafA1 did not increase the LC3-II/Total LC3 levels (**Figure S1c and d**) suggesting a defect in late stages of autophagy as expected. In CLN3-KO HeLa cells, we observed a higher level of LC3-II/Total LC3 levels compared to wild-type cells. Much like Rab7A-KO cells, the LC3-II/Total LC3 levels was increased in starvation conditions, and was not further increased by the BafA1 treatment (**Figure S1c and d**). Overall, our data from the CLN3-KO HeLa cells supports previously published results suggesting that CLN3 and Rab7A may function in a similar pathway to mediate starvation induced autophagy (Chandrachud et al., 2015).

To determine if CLN3 is required for the membrane recruitment of Rab7A, we performed a membrane isolation experiment as we have previously done (Mamo et al., 2012; Modica et al., 2017), in wild-type, CLN3-KO and CLN3-KO HeLa cells expressing FLAG-CLN3 (**Figure 2a**). Our membrane separation was successful, as the integral membrane protein Lamp2 was found in the pellet fraction (P), while the cytosolic protein tubulin was found in the supernatant fraction (S) (**Figure 2a**). Quantification of 5 independent experiments showed that Rab7A was not significantly displaced from membranes to the cytosolic fraction in CLN3-KO HeLa cells compared to wild-type HeLa cells, while the expression of Flag-CLN3 in CLN3-KO cells did not affect this phenotype (**Figure 2b**). The small GTPase Rab7A regulates the spatiotemporal recruitment of retromer (Rojas et al., 2008; Seaman et al., 2009), a protein complex required for efficient endosome-to-TGN traffic of cation-independent mannose 6-phosphate receptor (CI-MPR) and sortilin (Arighi et al., 2004; Canuel et al., 2008b; Seaman, 2004). In cells lacking retromer, these receptors do not efficiently recycle to the TGN, and are subsequently degraded in lysosomes (Arighi et al., 2004; Seaman, 2004). However, in Rab7A depleted cells, CI-MPR was not efficiently recycled to the TGN, but was not degraded (Rojas et al., 2008). We repeated the membrane assay as above, but we included Rab7A-KO HeLa cells a as control, as we previously demonstrated a reduction of retromer recruitment in Rab7A-KO HEK293 cells (Modica et al., 2017). Lamp2 and tubulin were used as markers of membrane (P) and cytosolic fractions (S) respectively (**Figure 2c**). Compared to wild-type HeLa cells, we observed a significant decrease in retromer recruitment (as detected by the retromer subunit Vps26A staining) in Rab7A-KO HeLa cells, but we did not observe any significant changes in CLN3-KO cells (**Figure 2c**). Quantification from 3 independent experiments found no changes in the membrane distribution of retromer in CLN3-KO and CLN3-KO cells expressing Flag-CLN3 compared to wild-type cells (**Figure 2d**). However, we found a 26% increase of retromer in the supernatant (cytosolic fraction) of Rab7A-KO cells compared to wild-type cells (**Figure 2d**), which is comparable to other previously published studies (Mamo et al., 2012; Seaman et al., 2009).

**Figure 2.**
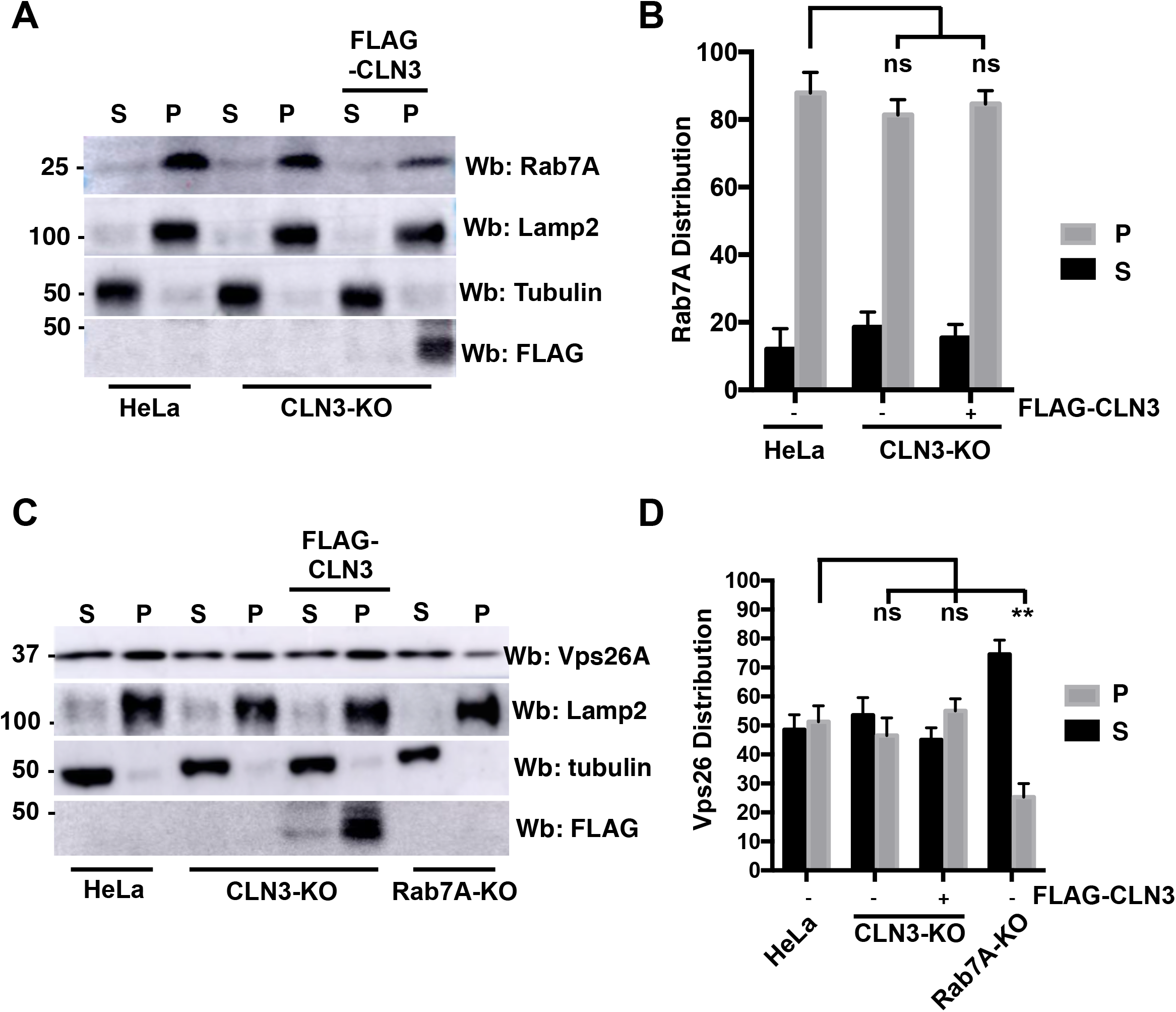
CLN3 is not required for the recruitment of Rab7A to membrane. Wild-type, CLN3-KO and FLAG-CLN3 expressing CLN3-KO HeLa cells were collected for membrane isolation assay. Samples were subjected to Western blot (Wb) with anti-Rab7A antibody, anti-Lamp2 antibody (a marker for the membrane fraction), and anti-tubulin antibody (a marker for the cytosolic fraction). Anti-FLAG staining is used to show the expression of wild-type CLN3 rescue. S, supernatant, P, pellet. (B) Quantification of 5 separate membrane isolation assay experiments. Data is represented as mean ± SD. ns, not significant; two-way ANOVA followed by Tukey’s post hoc test. (C) Wild-type, CLN3-KO, FLAG-CLN3 expressing CLN3-KO and Rab7A-KO HeLa cells were collected for a membrane isolation assay. After the isolation, samples were subjected to Western blot (Wb) with anti-Vps26A antibody (a retromer subunit), anti-Lamp2 antibody (a marker for the membrane fraction), and anti-tubulin antibody (a marker for the cytosolic fraction). Anti-FLAG staining is used to show the expression of wild-type CLN3. S, supernatant, P, pellet (D) Quantification of 5 separate membrane isolation assay experiments. Data is represented as mean ± SD. ns, not significant; **, P ≤ 0.01; two-way ANOVA followed by Tukey’s post hoc test.

### CLN3 is required for efficient retromer function

Although we found no changes in the membrane distribution of retromer in CLN3-KO cells, we wondered if we could observe other defects. Rab7A interacts with retromer, which is required for the spatiotemporal recruitment of the latter. We tested whether or not the interaction between Rab7A and retromer is affected in CLN3-KO cells. We generated BRET titration curves between RlucII-Rab7A and Vps26A-GFP10 in wild-type (blue curve) and CLN3-KO (red curve) HeLa cells (**Figure 3a**). Compared to wild-type HeLa cells, we found a 2 fold increase in the BRET_50_ values for Rab7A binding to Vps26A in CLN3-KO cells, suggesting a weaker interaction (**Figure 3b**). We previously demonstrated that the retromer/Rab7A interaction requires the palmitoylation of Rab7A (Modica et al., 2017). We tested whether or not the palmitoylation level of Rab7A was lower in CLN3-KO versus wild-type HeLa cells using Acyl-RAC, a technique to determine the palmitoylation status of a protein (**Figure 3c**). Quantification of 3 separate Acyl-RAC assays found no significant change in the level of palmitoylation of Rab7A in CLN3-KO cells compared to wild-type HeLa cells (**Figure 3d**).

**Figure 3.**
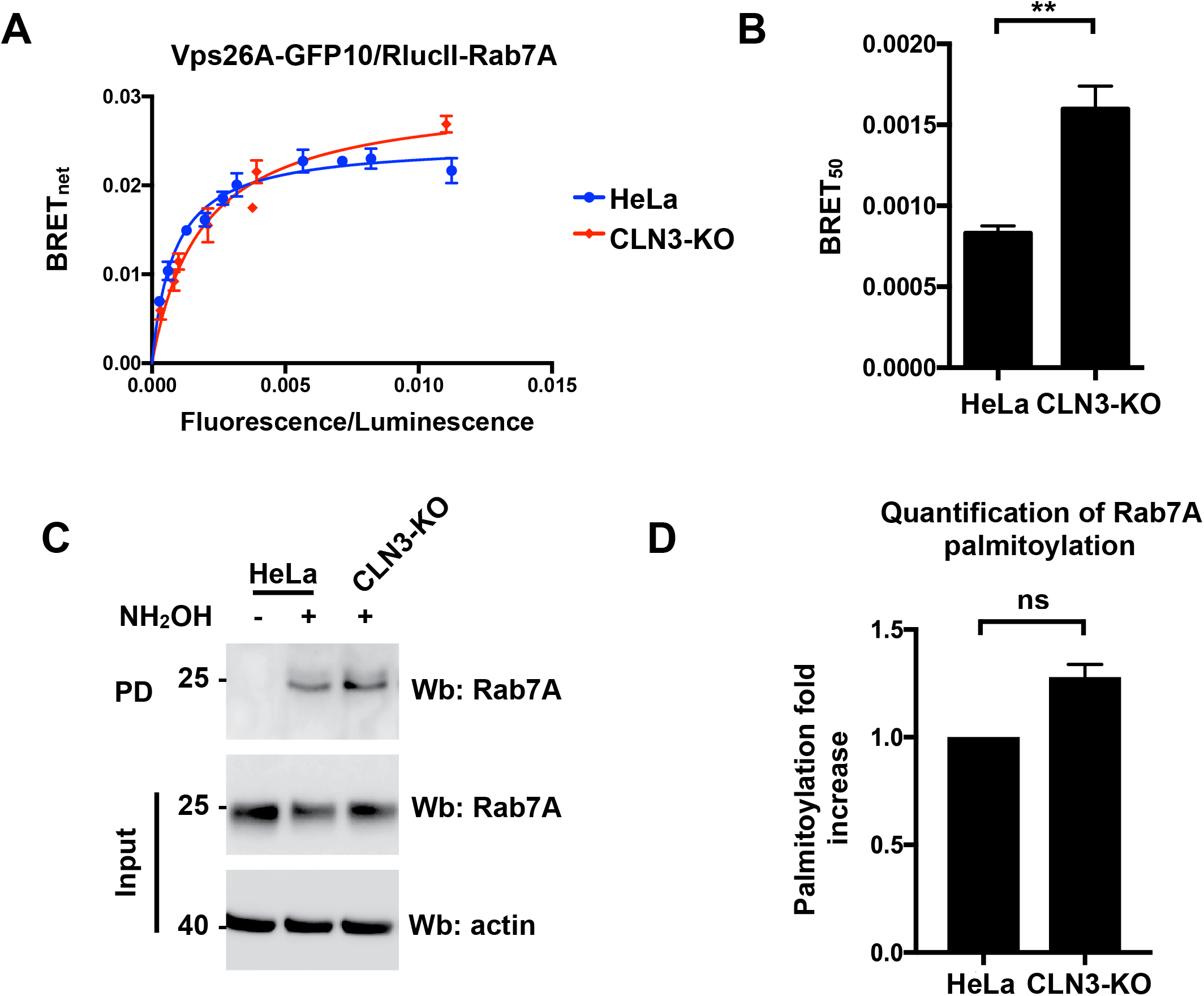
CLN3 is required for the efficient interaction of Rab7A with retromer. (A) Wild-type and CLN3-KO HeLa cells were transfected with a constant amount of RlucII-Rab7A and increasing amounts of Vps26A-GFP10 to generate BRET titration curves. 48 hours post-transfection BRET analysis was performed. BRET signals are plotted as a function of the ratio between the GFP10 fluorescence over RlucII luminescence. (B) BRET_50_ was extrapolated from 3 three independent experiments. Data is represented as mean ± SD. **, P ≤ 0.01; Student’s t-test (C) Wild-type and CLN3-KO HeLa cell lysates were collected and subjected to Acyl-RAC analysis and Wb with Rab7A antibody. NH_2_OH: Hydroxylamine. (D) Quantification of 3 separate Acyl-RAC assay experiments. Data is represented as mean ± SD. ns, not significant; Student’s t-test.

We next asked if CLN3 can interact with retromer. We generated BRET titration curves using RlucII-CLN3 and Vps26A-GFP10 (**Figure 4a, blue curve**). We identified an interaction between CLN3 and retromer. The CLN3^R334H^ (**Figure 4a, red curve**) and CLN3^E295K^ (**Figure 4a, black curve**) mutations had no impact on this interaction (**Figure 4b**). Interestingly, the CLN3^V330F^ (**Figure 4a, green curve**) increased the propensity of CLN3 to interact with retromer, while the CLN3^L101P^ (**Figure 4a, purple curve**) mutation significantly weakened the interaction (**Figure 4b**). CI-MPR and sortilin are known to interact with retromer, which is necessary for their endosome-to-TGN trafficking (Arighi et al., 2004; Canuel et al., 2008b). Using BRET, we confirmed this interaction using Vps26A tagged to nano-Luciferase (Vps26A-nLuc) and sortilin-YFP (**Figure 4c, black curve**). We next tested the interaction in CLN3-KO HeLa cells to determine if CLN3 played a role in this interaction (**Figure 4c, blue curve**). As expected, the sortilin/retromer interaction was significantly weaker in CLN3-KO and Rab7A-KO cells compared to wild-type HeLa cells (**Figure 4d**). As a control, we compared the interaction of sortilin and retromer in wild-type (**Figure 4e, black curve**) and Rab7A-KO (**Figure 4e, blue curve**) HeLa cells. As expected, we found a significantly weakened interaction between retromer and sortilin in Rab7A-KO cells compared to wild-type HeLa cells (**Figure 4f**).

**Figure 4.**
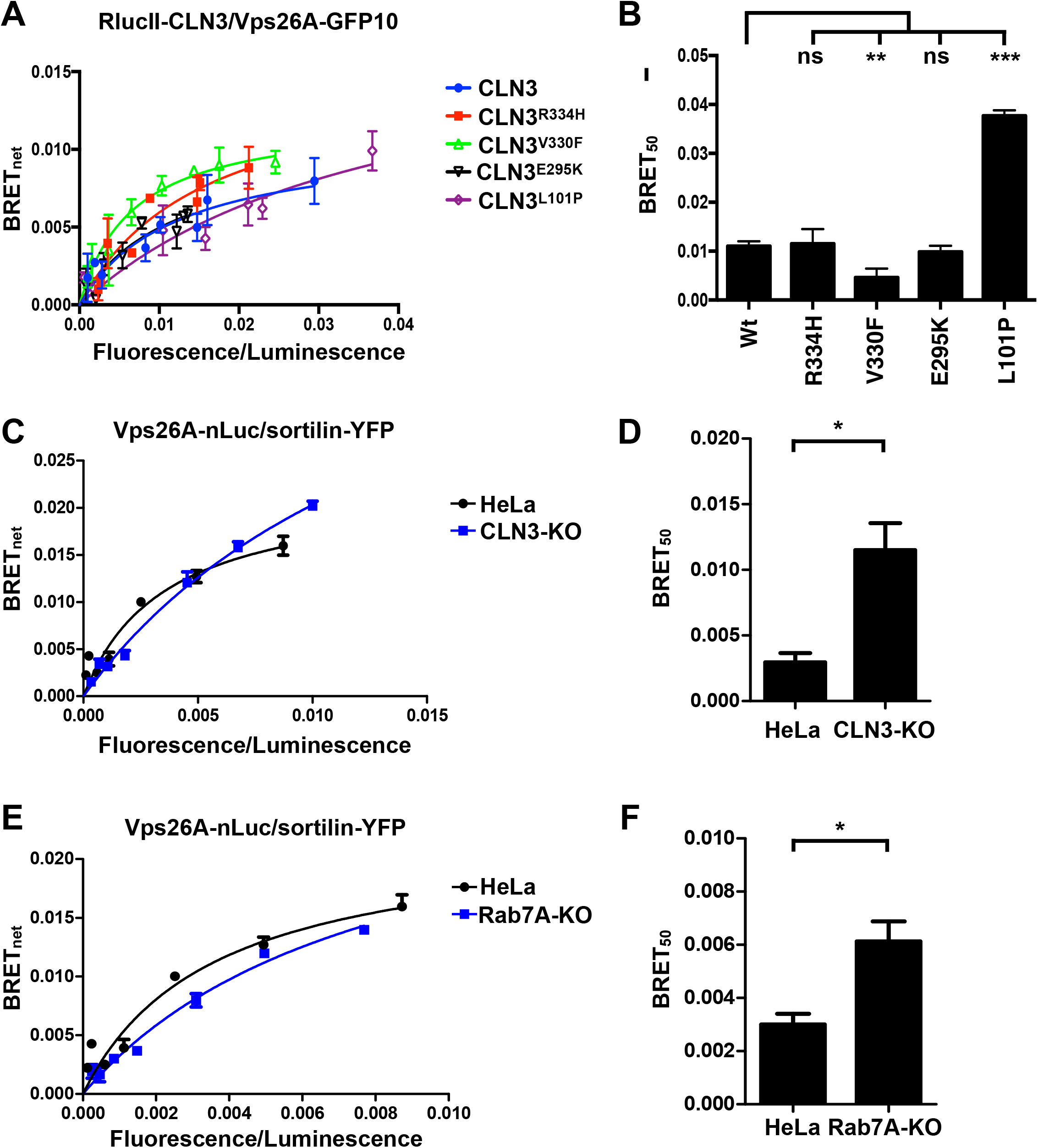
CLN3 interacts with retromer and modulates the sortilin/retromer interaction. (A) HeLa cells were transfected with a constant amount of RlucII-CLN3 or RlucII-CLN3 harbouring a disease-causing mutation and increasing amounts of Vps26A-GFP10 to generate BRET titration curves. 48 hours post-transfection, BRET analysis was performed. BRET signals are plotted as a function of the ratio between the GFP10 fluorescence over RlucII luminescence. (B) BRET_50_ was extrapolated from 3 independent experiments. Data is represented as mean ± SD. ns, not significant; **, P ≤ 0.01; ***, P ≤ 0.001; two-way ANOVA followed by Tukey’s post hoc test. (C) Wild-type and CLN3-KO HeLa cells were transfected with a constant amount of Vps26A-nLuc and increasing amounts of sortilin-YFP to generate BRET titration curves. 48 hours post-transfection BRET analysis was performed. BRET signals are plotted as a function of the ratio between the YFP fluorescence over nLuc luminescence. (D) BRET_50_ was extrapolated from 3 independent experiments. Data is represented as mean ± SD.*, P ≤ 0.05; two-way ANOVA followed by Tukey’s post hoc test. (E) Wild-type and Rab7A-KO HeLa cells were transfected with a constant amount of Vps26A-nLuc and increasing amounts of sortilin-YFP to generate BRET titration curves. 48 hours post-transfection BRET analysis was performed. BRET signals are plotted as a function of the ratio between the YFP fluorescence over nLuc luminescence. (F) BRET_50_ was extrapolated from 3 independent experiments. Data is represented as mean ± SD.*, P ≤ 0.05; two-way ANOVA followed by Tukey’s post hoc test.

### CLN3 regulates the stability of sortilin

Since we observed a weaker Rab7A/retromer interaction in CLN3-KO cells and retromer is required for the efficient endosome-to-TGN trafficking of CI-MPR and sortilin (Arighi et al., 2004; Canuel et al., 2008b), we wanted to determine if lysosomal sorting receptor recycling is regulated by CLN3. In cells depleted of retromer or retromer function, CI-MPR and sortilin are degraded in lysosomes rather than recycled back to the TGN (Arighi et al., 2004). We performed a cycloheximide chase experiment to determine receptor stability as we have previously done (Mamo et al., 2012; McCormick et al., 2008). 12 hours after wild-type, CLN3-KO and Rab7A-KO HeLa cells were seeded, the cells were incubated with serum free medium containing 50 μg/ml cycloheximide and collected after 0, 3, and 6 hours of incubation. Western blot (Wb) analysis shows decreased levels of sortilin and CI-MPR in CLN3-KO HeLa cell compared to Rab7A-KO and wild-type HeLa cells (**Figure S2a**). Transfecting FLAG-CLN3 in CLN3-KO rescued this phenotype (**Figure S2a**). Actin staining was used as a loading control, while FLAG staining was used to demonstrate the expression level of FLAG tagged constructs. Compared to wild-type cells which had 80% and 82% of sortilin remaining (**Figure 5a**) and 100% and 95% CI-MPR remaining (**Figure 5b**) at 3 and 6 hours, sortilin and CI-MPR were significantly degraded in CLN3-KO cells as only 30% and 19% of sortilin (**Figure 5a**) and 37% and 8% of CI-MPR (**Figure 5b**) remained after 3 and 6 hours. Expressing FLAG-CLN3 in CLN3-KO cells rescued recycling of sortilin (**Figure 5a**) and CI-MPR (**Figure 5b**) as they were no longer degraded and had protein levels remaining similar to wild-type HeLa cells with 74% and 77% and 94% and 95% remaining respectively at 3 and 6 hours. The amount of sortilin and CI-MPR remaining in Rab7A-KO cells (89% and 85% and 76% and 80% remaining respectively after 3 and 6 hours) was comparable to wild-type cells (**Figure 5a and b**). Next, we sought to determine the effects of CLN3 mutations known to cause human disease on the stability of these two cargo receptors. We expressed FLAG-CLN3^R334H^, FLAG-CLN3^V330F^, FLAG-CLN3^E295K^ and FLAG-CLN3^L101P^ in CLN3-KO HeLa cells (**Figure S2b**). 48 hours post-transfection, a cycloheximide chase as above was performed and total cell lysate was collected. Western blotting (Wb) with anti-FLAG antibody was used to indicate the expression level of the various expressed proteins, and antiactin staining was used as a loading control. Although the degradation of sortilin and CI-MPR were not as robust as in CLN3-KO cells, the receptors were significantly degraded in cells expressing CLN3^V330F^ (74% and 42% of sortilin and 28% and 25% of CI-MPR remaining after 3 and 6 hours), CLN3^E295K^ (51% and 51% of sortilin and 57% and 31% of CI-MPR remaining after 3 and 6 hours) and CLN3^L101P^ (35% and 44% of sortilin and 53% and 30% of CI-MPR remaining after 3 and 6 hours) expressing CLN3-KO HeLa cells, compared to wild-type cells (**Figure 5a and b**). Interestingly, CLN3^R334H^ was able to rescue the phenotype (82% and 70% remaining after 3 and 6 hours) for sortilin (**Figure 5a**), which is comparable to wild-type cells, but not for CI-MPR (55% and 45% remaining at 3 and 6 hours) (**Figure 5b**).

**Figure 5.**
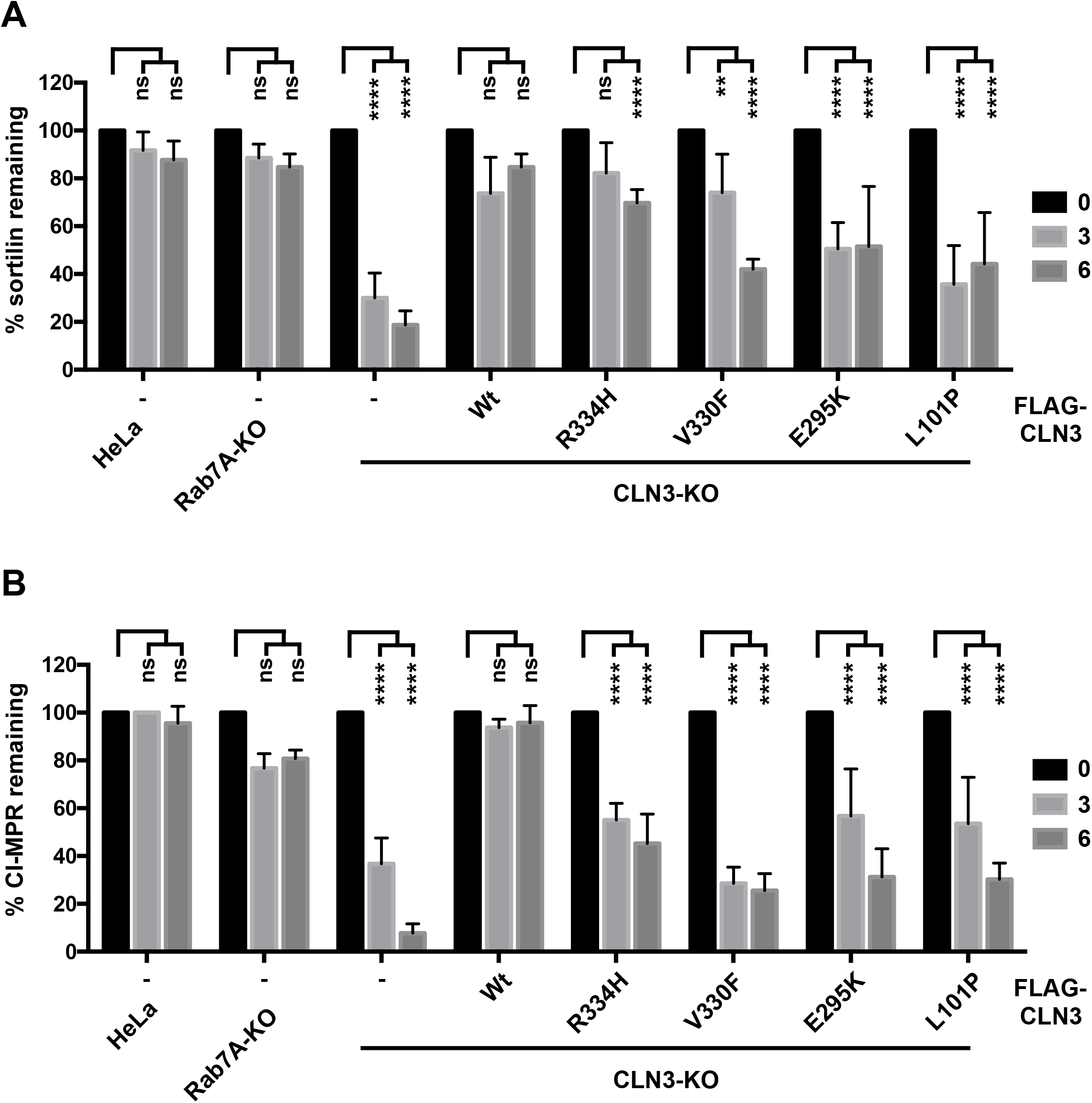
CLN3 regulates the stability of sortilin and CI-MPR. (A) Wild-type, Rab7A-KO, CLN3-KO or CLN3-KO HeLa cells expressing wild-type or mutant FLAG-CLN3 as indicated were incubated with 50 μg/ml cycloheximide in serum free media for 0, 3, or 6 hours. Quantification of 5 separate experiments (sample blots are shown in Figure S2) of sortilin remaining is plotted. Data is represented as mean ± SD. ns, not significant; ** ≤ 0.01, **** ≤ 0.0001; two-way ANOVA followed by Tukey’s post hoc test. (B) Wild-type, Rab7A-KO, CLN3-KO or CLN3-KO expressing wild-type or mutant FLAG-CLN3 as indicated were incubated with 50 μg/ml cycloheximide in serum free media for 0, 3, or 6 hours. Quantification of 5 separate experiments (sample blots are shown in Figure S2) of CI-MPR remaining at 6 hours in each group. Data is represented as mean ± SD. ns, not significant; **** ≤ 0.0001, two-way ANOVA followed by Tukey’s post hoc test.

### CLN3 is required for the efficient degradation of proteins following internalization

Upon EGF stimulation, EGFR is internalized and can be either recycled to the cell surface or degraded in lysosomes (Ceresa and Peterson, 2014). Rab7A is a key regulator of the degradative pathway mediating the later steps of this process (Ceresa and Bahr, 2006; Vanlandingham and Ceresa, 2009). At least two Rab7A effectors have been implicated in EGFR degradation, RILP and PLEKHM1. Indeed, depletion of either of these proteins results in significant delays in the degradation kinetics of EGFR (Marwaha et al., 2017; McEwan et al., 2015; Progida et al., 2007). In order to determine whether CLN3 modulates this Rab7A pathway, we investigated the degradation kinetics of EGFR in wild-type, CLN3-KO and Rab7A-KO HeLa cells. Wild-type, CLN3-KO and Rab7A-KO HeLa cells were serum starved for 1 hr in the presence of cycloheximide and then stimulated with 100 ng/ml of EGF in the presence of cycloheximide for the indicated periods of time. The level of endogenous EGFR was determined by Western blot and anti-actin staining was used as a control (**Figure 6a**). In wild-type cells, EGFR degradation was observed after 10 minutes and quantification of 5 independent experiments found substantial degradation at 15 (35% remaining), 30 (21% remaining) and 120 minutes (6% remaining) (**Figure 6b**). As expected, Rab7A-KO cells had significantly delayed degradation compared to wild-type cells at most indicated time points with 82% and 76% remaining at 10 and 15 minutes. However, in the Rab7A-KO HeLa cells, EGFR was degraded at 30 and 120 minutes. When we compared the degradation kinetics in CLN3-KO cells, we found significant delays at 10 (99% remaining), 15 (97% remaining) and 30 minutes (77% remaining) compared to wild-type cells, with no significant difference at 120 minutes (**Figure 6b**). The delayed degradation kinetics between Rab7A-KO and CLN3-KO was similar at 5, 10, 15 and 120 minutes, while the CLN3-KO cells contained significantly more EGFR at 30 minutes than Rab7A-KO HeLa cells.

**Figure 6.**
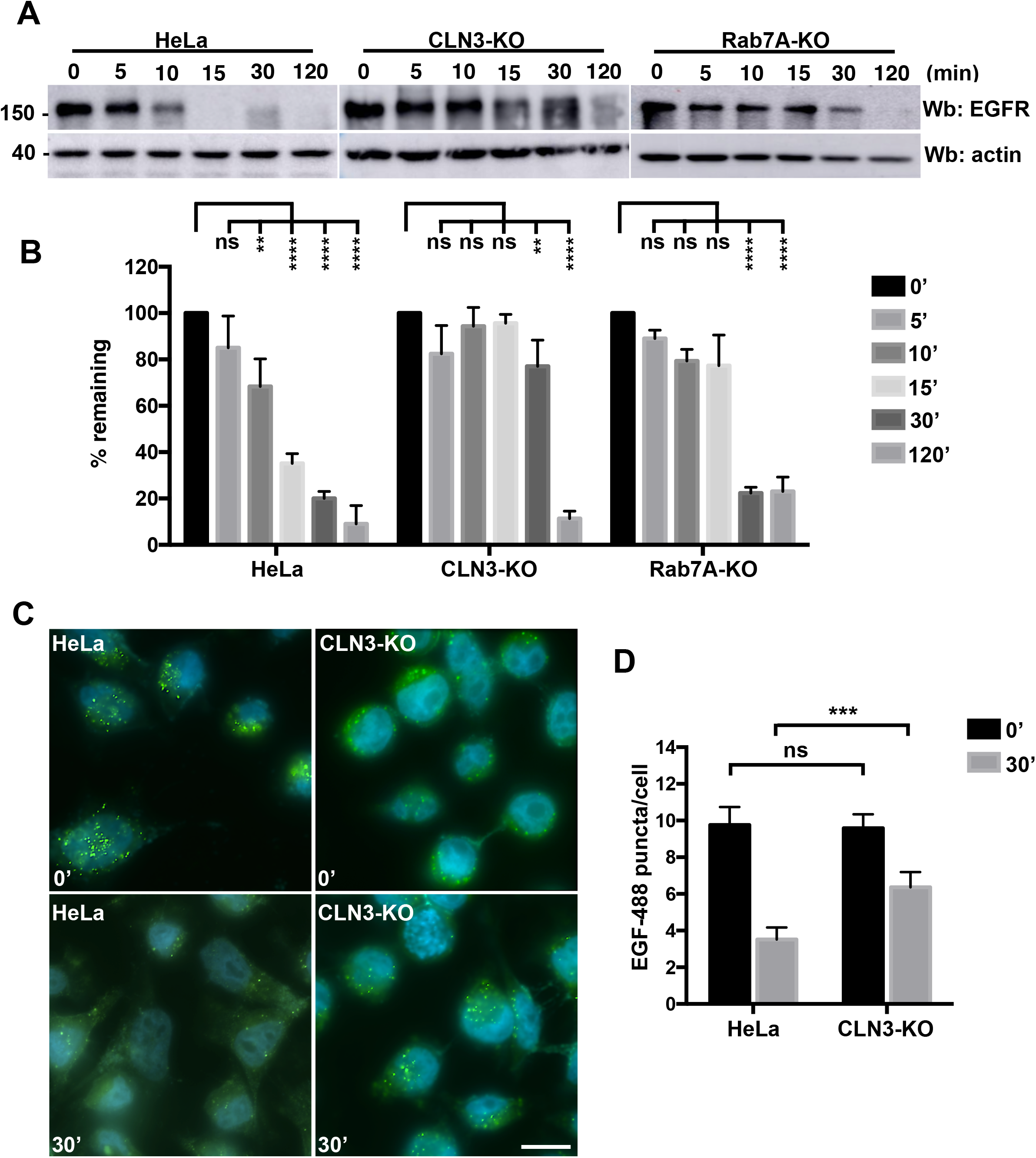
CLN3 is required for EGFR degradation. (A) Wild-type, CLN3-KO, and Rab7A-KO HeLa cells were incubated with 50 μg/ml cycloheximide for 1 hour and subsequently treated with 100 ng/ml EGF in Opti-MEM for 0, 5, 10, 15, 30, and 120 minutes. Whole cell lysate was then run on SDS-PAGE and a Western blot (Wb) was performed using anti-EGFR antibody. Anti-actin staining was used as a loading control. (B) Quantification of the remaining EGFR as detected in A was performed in 5 independent experiments. Data is represented as mean ± SD. ns, not significant; **, P ≤ 0.01, ***, P ≤ 0.001; two-way ANOVA followed by Tukey’s post hoc test. (C) Wild-type and CLN3-KO HeLa cells were grown on cover slips and incubated with 300 ng/ml EGF-488 for 0 or 30 minutes. The cells were fixed with 4% PFA for 12 minutes, followed by staining with DAPI to visualize the nucleus. Images were taken on a Zeiss Fluorescence microscope using a 63X objective. EGF-488 (green puncta) were counted manually in 50 cells per condition. (D) The results shown are the average number of puncta per cell per condition. Data is represented as mean ± SD. ns, not significant; ***, P ≤ 0.001; two-way ANOVA followed by Tukey’s post hoc test.

Next, we tested the degradation kinetics of Alexa-488 labeled EGF (EGF-488) using the same cell lines to confirm our EGFR degradation result. Following 2 hours of serum starvation, cells were incubated with 300 ng/ml EGF-488 for 30 minutes, washed and then chased for 0, or 30 minutes (**Figure 6c**). Images were acquired at random from the 4 different conditions and the number of EGF-488 punctate manually counted. 50 cells from each condition were counted. At time zero, both wild-type and CLN3-KO HeLa cells had comparable number of EGF-488 puncta (an average of 9.76 versus 9.58 respectively). After 30 minutes of chase, wild-type HeLa cells had an average of 3.52 puncta per cell, while CLN3-KO HeLa had on average 6.36 puncta per cell, an increase of over 1.8 times the number of puncta (**Figure 6d**). The delayed degradation kinetics observed in the CLN3-KO cells can be explained by decreased lysosomal function as a result of defective endosome-to-TGN trafficking, or it could be that EGF/EGFR do not reach the lysosomes efficiently due to lack of fusion.

In order to understand the mechanism behind the delayed EGFR degradation, we used BRET to determine if the Rab7A/RILP, Rab7A/PLEKHM1 and/or Rab7A/FYCO1 interactions were affected (**Figure 7a - f**). FYCO1 is a Rab7A effector required for anterograde traffic of vesicles (Pankiv et al., 2010), while RILP and PLEKHM1 are implicated in membrane fusion and degradation (Marwaha et al., 2017; McEwan et al., 2015; Progida et al., 2007). We found no significant change in the interaction between RILP and Rab7A (**Figure 7a and b**) or FYCO1 and Rab7A (**Figure 7c an d**) in either wild-type or CLN3-KO HeLa cells as shown by the BRET_50_ values. We did find a change in the interaction between PLEKHM1 and Rab7A, as the BRET_50_ value was 3.5 fold higher for the Rab7A/PLEKHM1 interaction in CLN3-KO cells compared to wild-type cells, suggesting a weaker interaction (**Figure 7e and f**).

**Figure 7.**
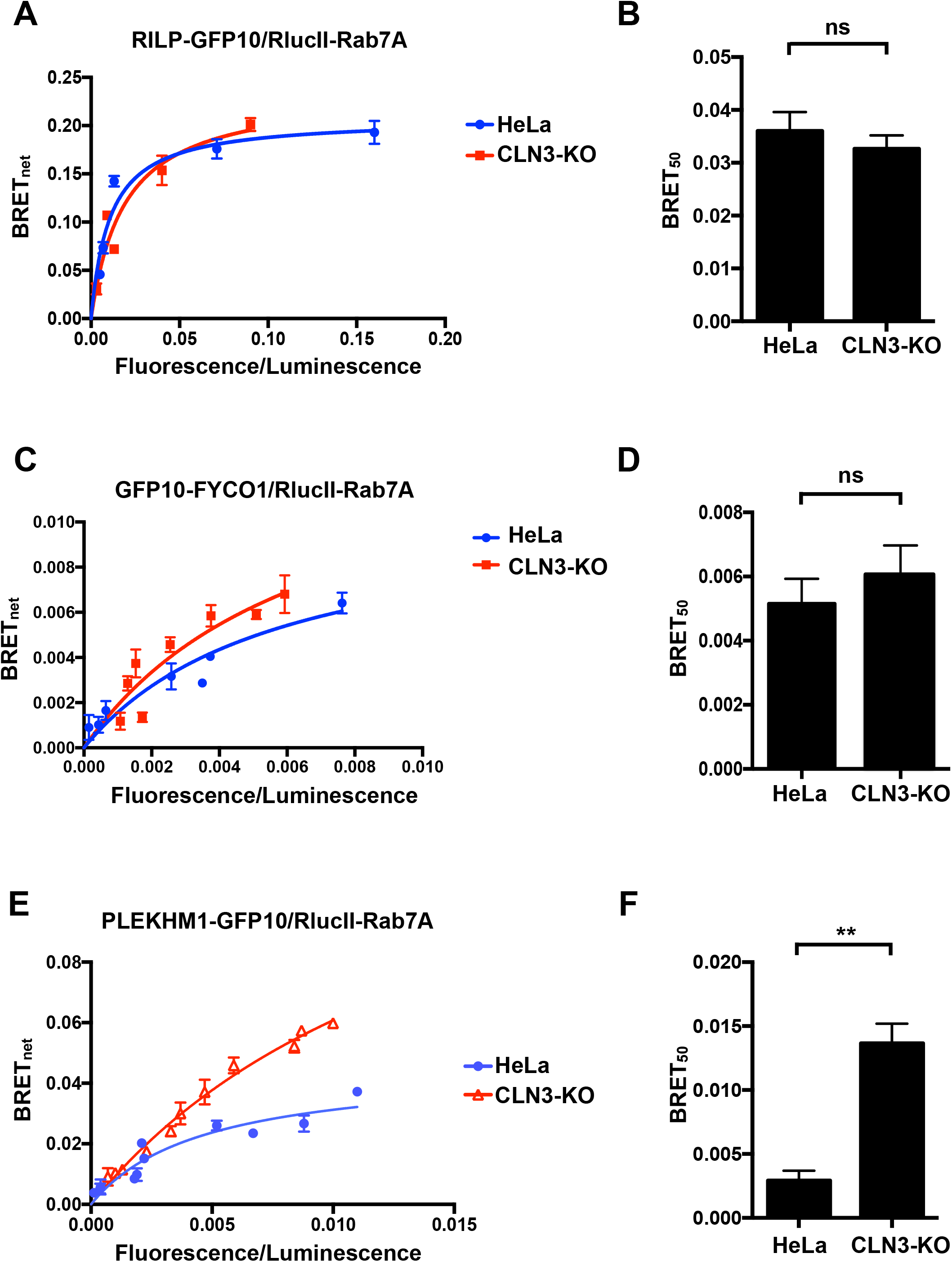
CLN3 regulates endocytic degradation by modulating the Rab7A/PLEKHM1 interaction. (A) Wild-type and CLN3-KO HeLa cells were transfected with a constant amount of RlucII-Rab7A and increasing amounts of RILP-GFP10 to generate BRET titration curves. BRET signals are plotted as a function of the ratio between the GFP10 fluorescence over RlucII luminescence. (B) BRET_50_ was extrapolated from 3 independent experiments. Data is represented as mean ± SD. ns, not significant; Student’s t-test (C) Wild-type and CLN3-KO HeLa cells were transfected with a constant amount of RlucII-Rab7A and increasing amounts of GFP10-FYCO1 to generate BRET titration curves. BRET signals are plotted as a function of the ratio between the GFP10 fluorescence over RlucII luminescence. (D) BRET_50_ was extrapolated from 3 independent experiments. Data is represented as mean ± SD. ns, not significant; Student’s t-test. (E) Wild-type and CLN3-KO HeLa cells were transfected with a constant amount of RlucII-Rab7A and increasing amounts of PLEKHM1-GFP10 to generate BRET titration curves. BRET signals are plotted as a function of the ratio between the GFP10 fluorescence over RlucII luminescence. (F) BRET_50_ was extrapolated from 3 independent experiments. Data is represented as mean ± SD. **, P ≤ 0.01; Student’s t-test.

## Discussion

CLN3 is an integral membrane protein localized to endosomes and lysosomes whose function has been implicated in intracellular trafficking and autophagy (Chandrachud et al., 2015; Metcalf et al., 2008; Oetjen et al., 2016). Previous work using *in vitro* techniques had demonstrated an interaction between CLN3 and the small GTPase Rab7A (Uusi-Rauva et al., 2012). We confirmed this interaction using bioluminescence resonance energy transfer (BRET). Furthermore, we showed that two CLN3 mutations, CLN3^R334H^ and CLN3^V330F^, increased this interaction. This suggests that these two mutations could retain Rab7A on membranes longer or prevent its efficient cycling, a process required for optimal function. Interestingly two other point mutation, CLN3^E295K^ and CLN3^L101P^, had no effect on this interaction. However, these two mutations weakened the CLN3/sortilin interaction. Combined with CLN3 being required for both the Rab7A/retromer and retromer/sortilin interaction, our data reveals a role for CLN3 in modulating retromer function. Ablation of CLN3 results in the lysosomal degradation of sortilin and CI-MPR, which is most likely due to deficient endosome-to-TGN trafficking of sortilin and CI-MPR. A previous publication demonstrated defects in the trafficking and processing of the lysosomal enzyme cathepsin D in CLN3^Δex7-8^/CLN3^Δex7-8^ cells (Fossale et al., 2004). It is well established that defects in retromer function or sortilin affect cathepsin D sorting and processing (Arighi et al., 2004; Canuel et al., 2008a). As such, our results provide a molecular explanation to those previous observations.

Rab7A also plays a crucial role in the degradation of endocytic cargo such as EGFR (Vanlandingham and Ceresa, 2009). We found significant delays in the degradation of both EGFR and EGF in CLN3-KO cells. This could be explained in two ways. First, CLN3 is involved in the trafficking of the lysosomal sorting receptor sortilin. Defects in trafficking of this protein, or defects in retromer function have been shown to have a significant impact on lysosome function (Arighi et al., 2004; Lefrancois et al., 2003; Seaman, 2004). Second, Rab7A is required for the fusion of endosomes and lysosomes, a process requiring RILP and PLEKHM1 (Marwaha et al., 2017; McEwan et al., 2015; Progida et al., 2007). Although we found no change in the Rab7A/ RILP interaction in CLN3-KO cells, we found a significant decrease in the Rab7A/PLEKHM1 interaction. This suggest defective fusion events. Combined with decreased lysosomal function, this would explain the significant delay in EGF and EGFR degradation. The Akt-mTOR pathway is upregulated in CLN3 patient fibroblasts (Vidal-Donet et al., 2013). EGFR is known to activate several signalling pathways, including Akt (Mattoon et al., 2004). Disregulated trafficking and degradation can lead to increased signalling (Sorkin and von Zastrow, 2009). Our results could at least partially explain the increased Akt-mTOR pathway found in CLN3 patient fibroblasts.

Autophagy is affected in CLN3 deficient cells (Chandrachud et al., 2015; Chang et al., 2011; Vidal-Donet et al., 2013). Our CLN3-KO HeLa cells also show defective autophagy, similar to the defects observed in Rab7A-KO. This suggests defects late in the autophagy pathway, possibly at the fusion step with lysosomes. PLEKHM1, among other factors, plays a critical role in modulating autophagosome fusion with lysosomes (McEwan et al., 2015). Defects in this fusion machinery leads to defective autophagy, but also to slowed EGFR degradation kinetics. We propose that defects in autophagy observed in CLN3 disease patients could be due to defects in PLEKHM1 function, along with defects in lysosome function due to decreased sorting of sortilin and CI-MPR.

In conclusion, our results point to a role of CLN3 in the regulation of Rab7A function. We have previously shown a similar role for CLN5, a soluble protein found within the lumen of endosomes and lysosomes (Mamo et al., 2012). As CLN3 and CLN5 are known to interact, we propose that the two proteins function as a complex to regulate Rab7A and retromer function, demonstrating, at least partially, the molecular mechanisms deficient in CLN3 disease.

## Materials and Methods

### Plasmids and Mutagenesis

RlucII-Rab1a was generated by amplifying Rab1a cDNA from myc-Rab1a and cloned into the EcoRV/XhoI sites of pcDNA3.1 Hygro (+) RlucII-GFP10-st2 plasmid. RlucII-CLN3 and GFP10-CLN3 were generated by amplifying CLN3 cDNA from FLAG-CLN3 and cloned into the EcoRV/ XhoI and EcoRV/XbaI sites of pcDNA3.1Hygro(+)RlucII-GFP10-st2 and pcDNA3.1Hygro(+) GFP10-RlucII-st2 plasmids, respectively. To generate Vps26A-nano-Luc, Vps26A cDNA was obtained by PCR from Vps26A-YFP plasmid. pnLuc-N1 was generated by replacing the eYFP of pEYFP-N1 with nano-Luc (a generous gift from Regis Grailhe, Pasteur Institute Korea). Cloning was done after digestion of pnLuc-N1 plasmid with XhoI/HindIII restriction enzymes. PLEKHM1-GFP10 was engineered by inserting PLEKHM1 into the NheI/EcoRV sites pcDNA3.1 Hygro (+) GFP10-RLucII-st2. To generate GFP10-FYCOI, cDNA of FYCOI was obtained by PCR from mCherry-FYCOI and cloned into the KpnI/XbaI sites of pcDNA3.1 Hygro (+)-GFP10-RLucII. The various mutants were engineered using site-directed mutagenesis from the previously described FLAG-CLN3, GFP10-CLN3, RlucII-CLN3 constructs. For a detailed list of plasmids used in this study, please refer to Table S1.

### Cell culture and transient transfections

HeLa cells were cultured in Dulbecco’s modified Eagle’s medium (DMEM) supplemented with 2mM L-Glutamine, 100 U/ml penicillin, 100g/ml streptomycin and 10% FBS (Thermo Fisher Scientic, Burlington, ON) at 37 °C in a humidified chamber at 95% air and 5% CO_2_. Cells were seeded at a density of 2 × 10^5^/well for 12 well plates and 5 × 10^5^/well for 6 well plates 24 hours prior to transfection. 1 μg/well for 12 well plates and 2 μg/well for 6 well plates of total DNA were used for transient transfection using linear 25 kDa polyethyleneimine (3 μg PEI/μg DNA).

### CRISPR/Cas9 Editing

In order to generate CLN3 knockout cells, a guide RNA (gRNA) corresponding to the first exon of CLN3 was designed (CACCGCGGCGCTTTTCGGATTCCGA) and cloned into pX330-U6 expressing a humanized Cas 9 (Cong et al., 2013). HeLa cells were co-transfected with pX330-U6-gRNA-CLN3 and pcDNA3.1-zeocin (Invitrogen). 24 hours after transfection cells were selected with zeocin (250μg/ml) for 5 days. Cells were cultured for 2 weeks, single clones were isolated and genomic DNA extracted. DNA sequencing was performed to identify CLN3 knockout cells carrying a specific indel mutation. HeLa cells were transfected with an all-in-one CRISPR/Cas9 plasmid for Rab7A (plasmid number HTN218819, Genecopoeia, Rockville, MD). 72 hrs post-transfection, the cells were treated with 1mg/ml Geneticin (Thermo Fisher Scientific) for 1 week. Limited dilution was performed to isolate single cells which were allowed to grow for 2 weeks. Western blotting was used to identify Rab7A-KO cells. For a list of antibodies used in this study, please refer to Table S2.

### BRET titration experiments

HeLa cells were seeded in 12-well plates and transfected with the indicated plasmids. 48 hour post transfection, cells were washed in PBS, detached with 5mM EDTA in PBS and collected in 500 μl of PBS. Cells were transferred to opaque 96-well plates (VWR Canada, Mississauga, ON) in triplicates. Total fluorescence was first measured with the infinite M1000 Pro plate reader (Tecan Group Ltd., Mannedorf, Switzerland) with the excitation and emission set at 400 nm and 510 nm respectively for BRET^2^ and 500 nm and 530 nm for BRET^1^. The BRET^2^ substrate coelenterazine 400a and BRET^1^ substrate h-coelenterazine were then added to all wells (5 μM final concentration) and the BRET^2^ and BRET^1^ signals were measured 3 min later. The BRET signals were calculated as a ratio of the light emitted at 525 ± 15 nm over the light emitted at 410 ± 40 nm. The BRETnet signals were calculated as the difference between the BRET signal in cells expressing both fluorescence and luminescence constructs and the BRET signal from cells where only the luminescence fused construct was expressed.

### Western blotting

Cell were detached using 5mM EDTA in PBS, washed in 1X PBS and collected by centrifugation. TNE buffer (150 mM NaCl, 50 mM Tris, pH 7.5, 2 mM EDTA, 0.5% Triton X-100 and protease inhibitor cocktail) was used to lyse cells by incubating them for 30 minutes on ice. Lysates were centrifuged at high speed for 10 minutes and the supernatants (cell lysate) were collected to be analyzed by Western blotting. Samples were mixed with sample buffer 3X to obtain a final concentration of 1X (62.5 mM Tris-HCl pH 6.5, 2.5% SDS, 10% glycerol, 0.01% bromophenol blue). Prior to electrophoresis samples were incubated at 95°C for 5 minutes and resolved on SDS-PAGE followed by wet-transfer to nitrocellulose membranes. Detection was done by immunoblotting using the indicated antibodies.

### Membrane seperation assay

24 hours post-transfection, cells were collected in 5mM EDTA in PBS. The cells were subsequently snap frozen in liquid nitrogen and allowed to thaw at RT for 5min. The cells were resuspended in Buffer 1 (0.1 M Mes-NaOH pH 6.5, 1 mM MgAc, 0.5 mM EGTA, 200 M sodium orthovanadate, 0.2 M sucrose) and centrifuged at 10,000 g for 5 minutes at 4°C. The supernatant containing the cytosolic proteins (S, soluble fraction) was collected. The remaining pellet was resuspended in Buffer 2 (50 mM Tris, 150 mM NaCl, 1 mM EDTA, 0.1% SDS, 1% Triton X-100) and centrifuged at 10,000 g for 5 minutes at 4°C. Samples were loaded into SDS-PAGE gels in equal volumes. Fiji was used to quantify the intensity of the bands (Schindelin et al., 2012). The intensity of each fraction was calculated and divided by the total intensity to determine the distribution of proteins.

### Cycloheximide chase

Wild-type, CLN3-KO, Rab7A-KO HeLa cells were seeded in 6-well plates the day prior to transfection. 500 ng of FLAG fused wild-type and mutation harbouring CLN3 was transfected into CLN3-KO HeLa cells. 48 hours after transfection, cells were treated with 50 μg/ml of cycloheximide in Opti-MEM. Lysates were collected at 0, 3 or 6 hour-time points and run on 10% SDS-PAGE gels.

### EGFR degradation assay

Wild-type, CLN3-KO and Rab7A-KO cells were seeded in 6-well plates the day before the assay. In order to prevent de novo synthesis EGFR during EGF stimulation, cells were treated with 50 μg/ml cycloheximide in Opti-MEM for 1 hour. EGF stimulation was performed with 100 ng/ml EGF in Opti-MEM containing 50 μg/ml of cycloheximide. Cell lysates were collected as indicated above and Western blotting was performed. EGFR levels were measured using Fiji.

### EGF-488 pulse-chase

Wild-type and CLN3-KO HeLa cells were seeded on coverslips the day before the experiment. Cells were serum-starved in Opti-Mem for 1 h followed by a 30-min pulse of 300 ng/ml of EGF-488. Cells were then washed with PBS, fixed in 4% paraformaldehyde at 0 and 30 min and mounted onto slide using Fluoromount-G. The coverslips were sealed using nail polish. Cells were imaged using fluorescence microscope. The number of puncta per cell was counted manually (50 cells per condition for each time point).

### Autophagic flux

Wild-type, CLN3-KO, Rab7A-KO HeLa cells were seeded in 6-well plates the day before autophagy induction. Cells are starved with EBSS for 4 hours to induce autophagy. 100 nM of Bafilomycin A1 (BafA1) was used to inhibit lysosomal function and therefore inhibit autophagy. Lysates were run on SDS-PAGE and blotted with anti-LC3 and anti-actin antibodies. Band intensity was determined using Fiji, and quantification was performed as LC3-II over LC3-I + LC3-II.

### Statistics

Statistical analysis was performed using GraphPad Prism Version 7. The statistical tests used are described in the corresponding figure legends.

## Acknowledgements

This work was supported by grants from the Joint Programme in Neurodegenerative Diseases Grant/Canadian Institutes for Health Research (ENG-387575) and the Canadian Foundation for Innovation (35258) to SL. Parts of this work were supported by grants from the National Contest for Life (NCL) Foundation, Germany to GH and by the Deutsche Forschungsgemeinschaft to GH. GM is supported by a scholarship from Fond de recherche du Quebec – Santé.

## Author Contributions

Conceptualization: S.Y., S.L.; Methodology: S.Y., G.M., E.S., S.L., A.K., G.H.; Formal analysis: G. M., E.S., S.L., A.K., G.H.; Investigation: O.S., E.S., A.V.; Writing - original draft: S.Y., S.L.; Writing – review & editing: S.Y., G.M., S.L., G.H.; Supervision: S.L., G.H.; Funding acquisition: S.L., G.H.

---

**Figure S1.**
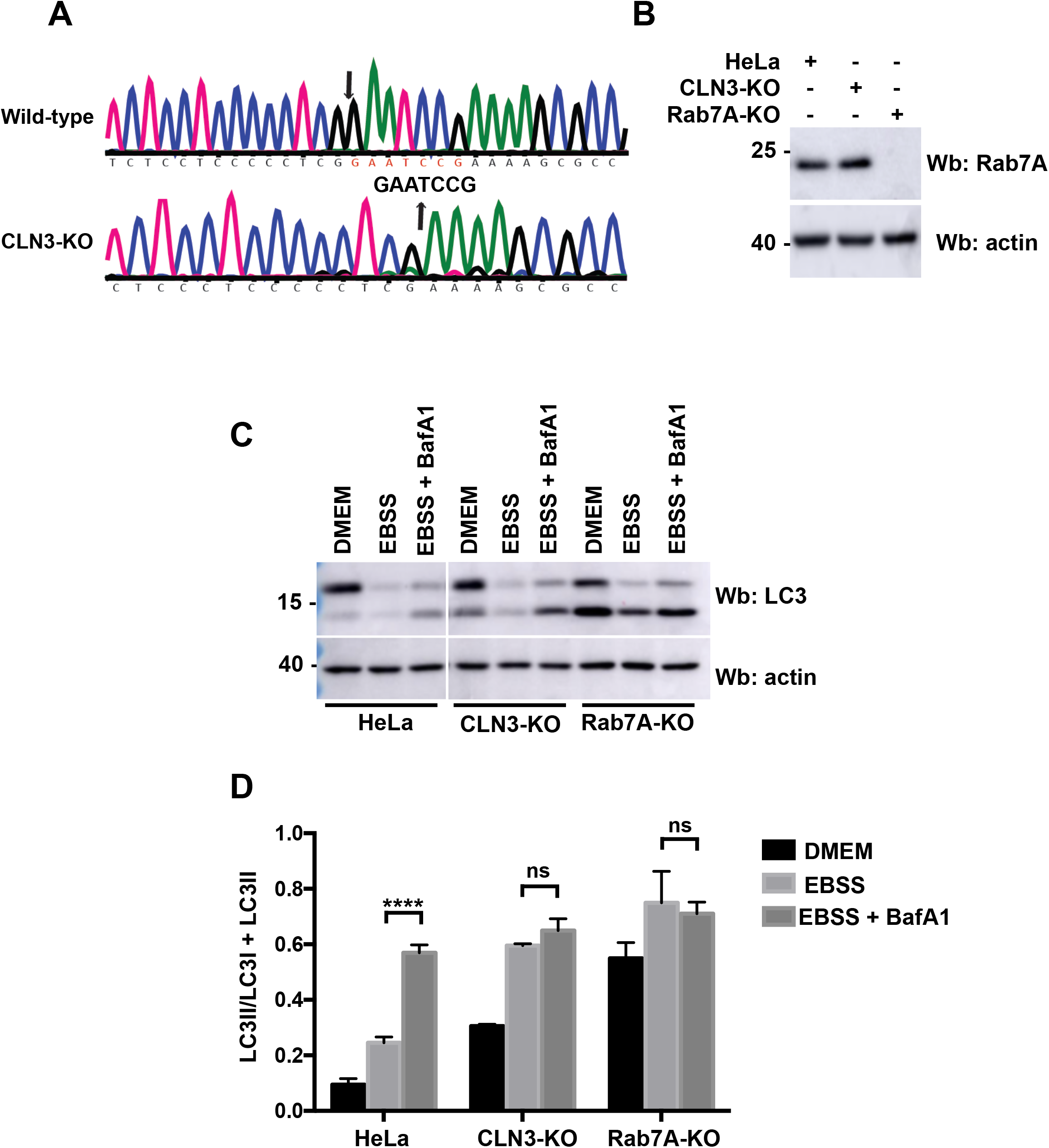
Engineering of CLN3 and Rab7A knockout cells. (A) DNA sequencing of genomic DNA of wild type (upper panel) and CLN3 knockout HeLa cells demonstrates the deletion of 7 base pairs in the CLN3 gene from position 35 to 41 after the start codon. Note that the reverse complementary sequence is shown. The deletion causes a frameshift, changes the encoded amino acid sequence starting at position 13 (D^13^ to R) and leads to a premature stop codon after amino acid 52. Arrow indicates the position of the Indel mutation. (B) Total cell lysate from wild-type, CLN3-KO and Rab7A-KO cells was run on an SDS-PAGE and Western blotting (Wb) was performed using anti-Rab7A and anti-actin antibodies. (C) Wild-type, CLN3-KO and Rab7A-KO HeLa cells were cultured in DMEM, EBSS, EBSS + Bafilomycin A1 (BafA1) for 4 hours. Total cells lysates were run on a SDS-PAGE and Western blotting (Wb) was performed with anti-LC3 and anti-actin antibodies. The amount of LC3-II was calculated as a ratio between the amount of LC3-II over total LC3 (LC3-I +LC3-II). (D) The results shown are representative of 5 independent experiments. Data is represented as mean ± SD. ns, not significant; ****, P ≤ 0.0001; two-way ANOVA followed by Tukey’s post hoc test.

**Figure S2.**
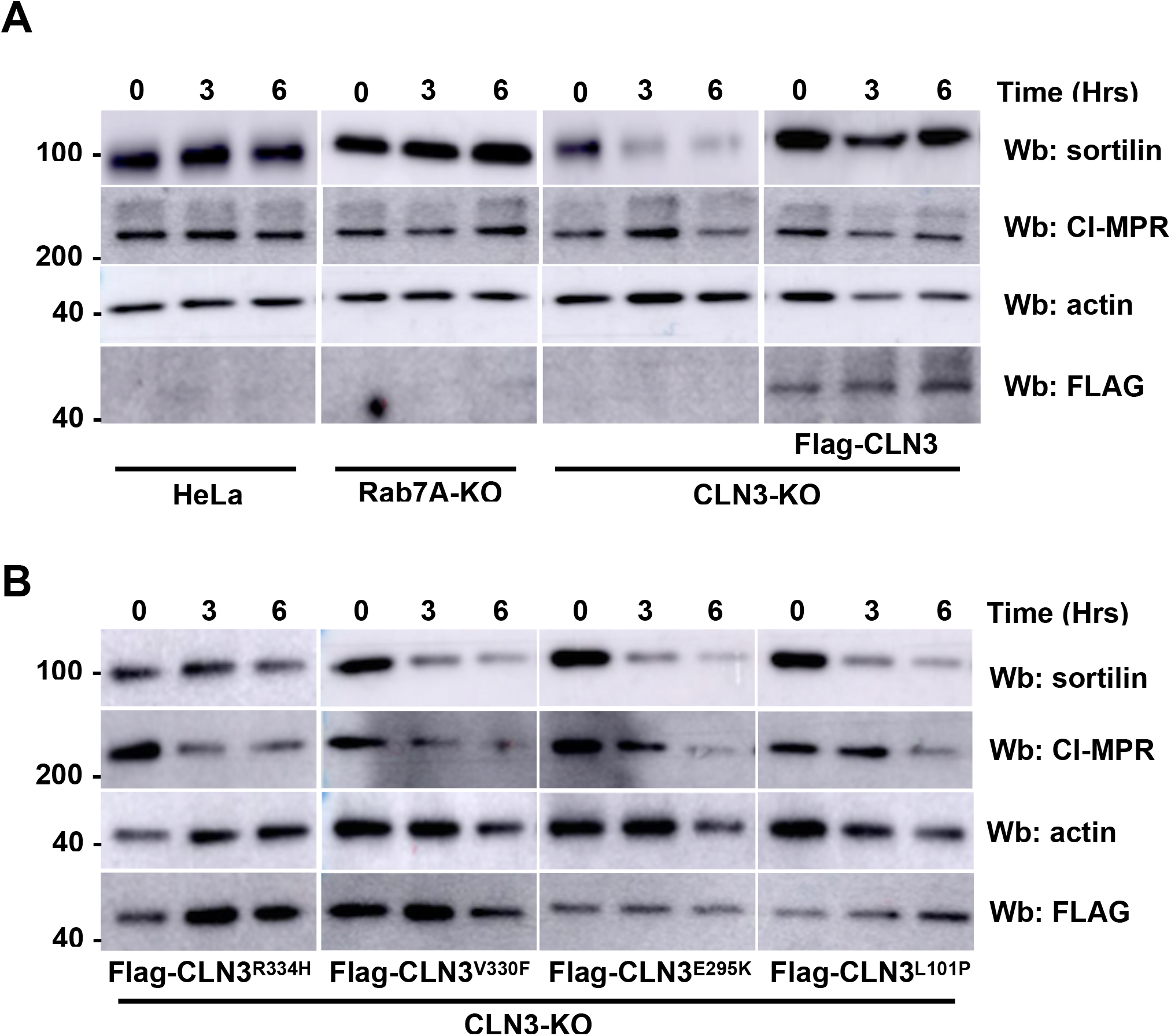
Stability of sortilin and CI-MPR. (A) Wild-type, Rab7A-KO, CLN3-KO and CLN3-KO expressing FLAG-CLN3 HeLa cells were incubated with 50 μg/ml of cycloheximide for the indicated times. Total cell lysate was run on SDS-PAGE and Western blotting was performed using anti-sortilin, anti-CI-MPR, anti-actin and anti-FLAG antibodies. (B) CLN3-KO HeLa cells expressing wild-type and mutant FLAG-CLN3 as indicated were incubated with 50 μg/ml of cycloheximide for the indicated times. Total cell lysate was run on SDS-PAGE and Western blotting was performed using anti-sortilin, anti-CI-MPR, anti-actin and anti-FLAG antibodies.

**Table S1.**
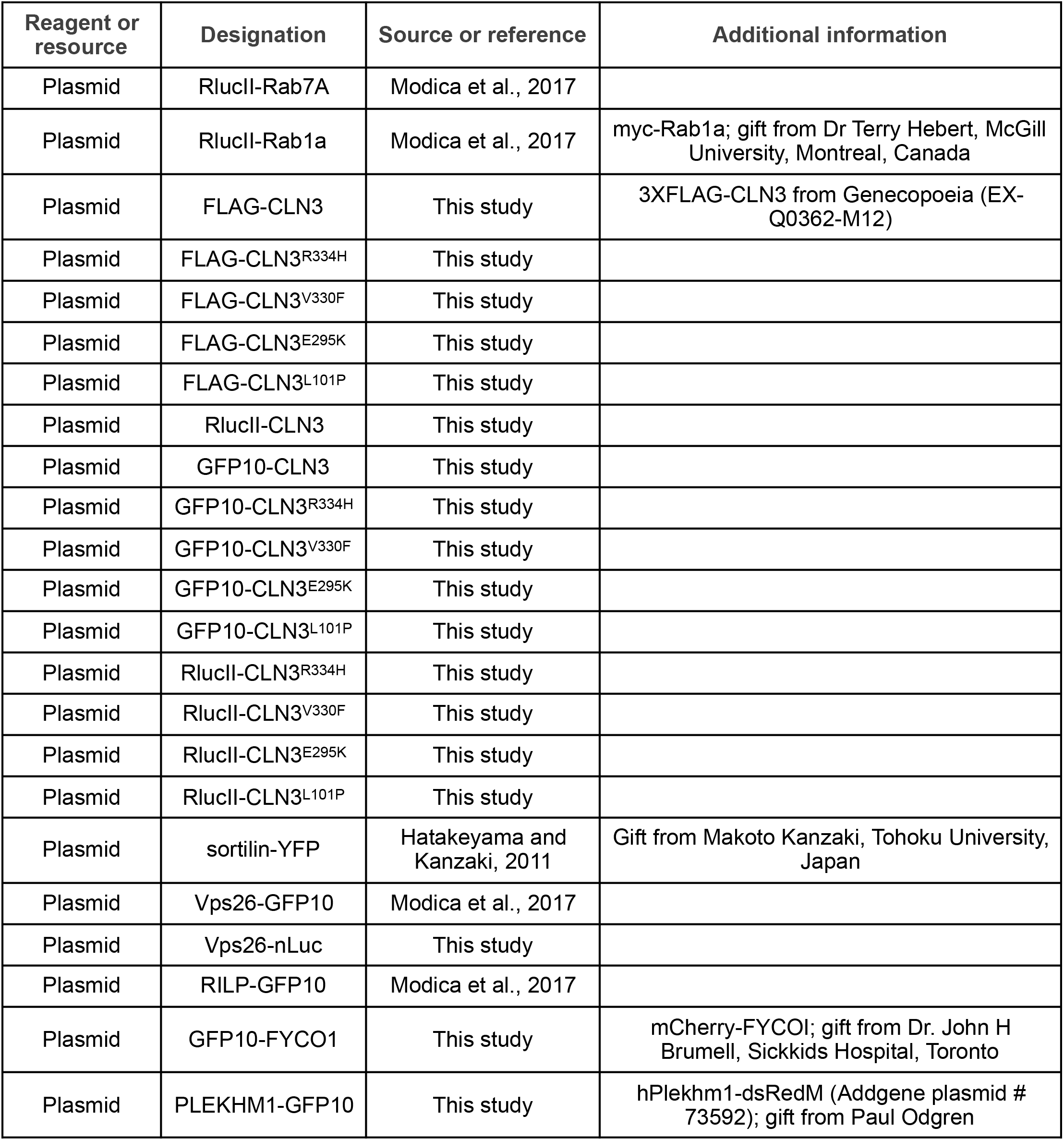
List of expression plasmids used in this study.

| Reagent or resource | Designation | Source or reference | Additional information |
| --- | --- | --- | --- |
| Plasmid | RlucII-Rab7A | Modica et al., 2017 |  |
| Plasmid | RlucII-Rab1a | Modica et al., 2017 | myc-Rab1a; gift from Dr Terry Hebert, McGill University, Montreal, Canada |
| Plasmid | FLAG-CLN3 | This study | 3XFLAG-CLN3 from Genecopoeia (EX-Q0362-M12) |
| Plasmid | FLAG-CLN3<sup>R334H</sup> | This study |  |
| Plasmid | FLAG-CLN3<sup>V330F</sup> | This study |  |
| Plasmid | FLAG-CLN3<sup>E295K</sup> | This study |  |
| Plasmid | FLAG-CLN3<sup>L101P</sup> | This study |  |
| Plasmid | RlucII-CLN3 | This study |  |
| Plasmid | GFP10-CLN3 | This study |  |
| Plasmid | GFP10-CLN3<sup>R334H</sup> | This study |  |
| Plasmid | GFP10-CLN3<sup>V330F</sup> | This study |  |
| Plasmid | GFP10-CLN3<sup>E295K</sup> | This study |  |
| Plasmid | GFP10-CLN3<sup>L101P</sup> | This study |  |
| Plasmid | RlucII-CLN3<sup>R334H</sup> | This study |  |
| Plasmid | RlucII-CLN3<sup>V330F</sup> | This study |  |
| Plasmid | RlucII-CLN3<sup>E295K</sup> | This study |  |
| Plasmid | RlucII-CLN3<sup>L101P</sup> | This study |  |
| Plasmid | sortilin-YFP | Hatakeyama and Kanzaki, 2011 | Gift from Makoto Kanzaki, Tohoku University, Japan |
| Plasmid | Vps26-GFP10 | Modica et al., 2017 |  |
| Plasmid | Vps26-nLuc | This study |  |
| Plasmid | RILP-GFP10 | Modica et al., 2017 |  |
| Plasmid | GFP10-FYCO1 | This study | mCherry-FYCO1; gift from Dr. John H Brumell, Sickkids Hospital, Toronto |
| Plasmid | PLEKHM1-GFP10 | This study | hPlekM1-dsRedM (Addgene plasmid # 73592); gift from Paul Odgren |

**Table S2.**
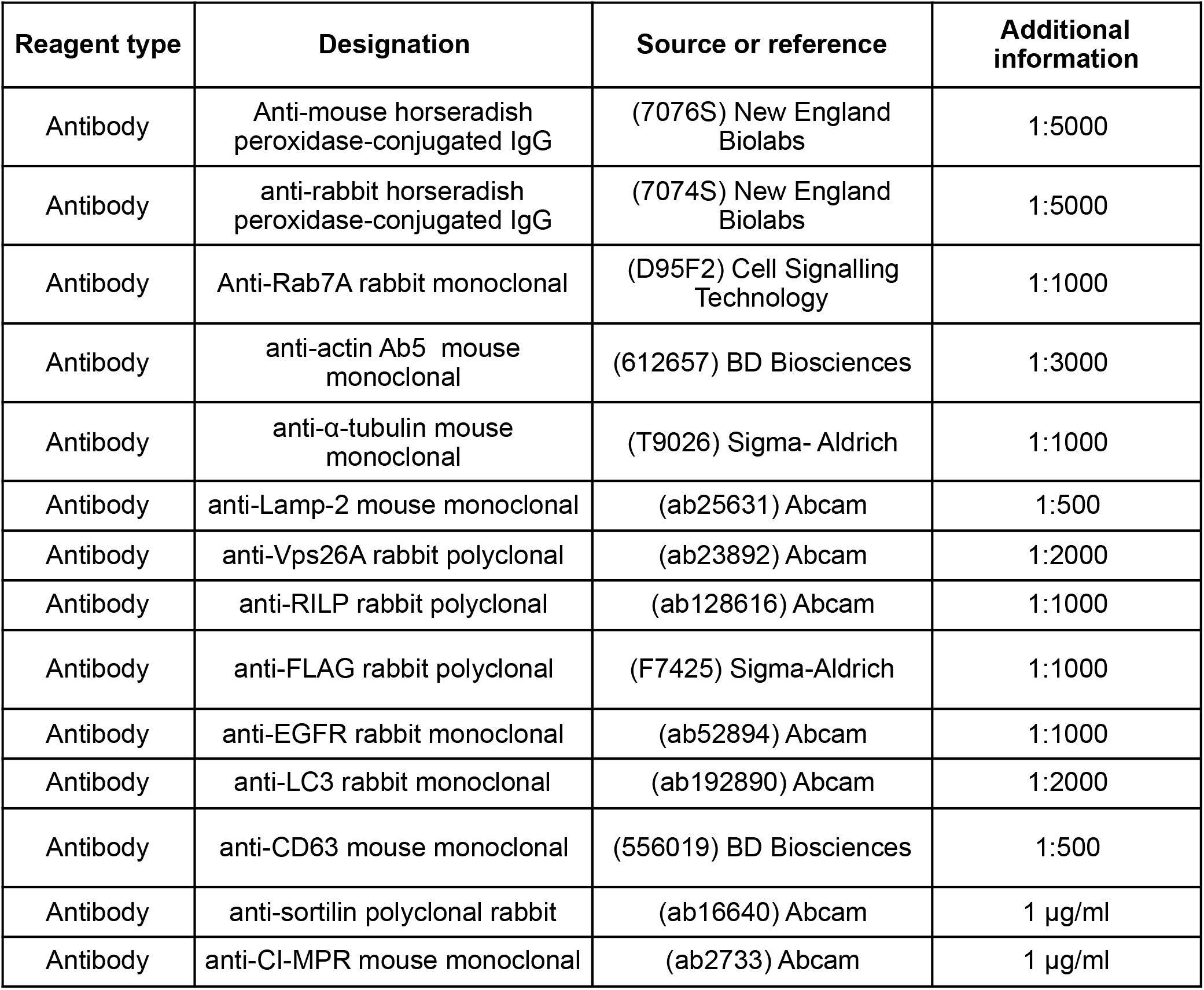
List of antibodies used in this study.

| Reagent type | Designation | Source or reference | Additional information |
| --- | --- | --- | --- |
| Antibody | Anti-mouse horseradish peroxidase-conjugated IgG | (7076S) New England Biolabs | 1:5000 |
| Antibody | anti-rabbit horseradish peroxidase-conjugated IgG | (7074S) New England Biolabs | 1:5000 |
| Antibody | Anti-Rab7A rabbit monoclonal | (D95F2) Cell Signalling Technology | 1:1000 |
| Antibody | anti-actin Ab5 mouse monoclonal | (612657) BD Biosciences | 1:3000 |
| Antibody | anti-α-tubulin mouse monoclonal | (T9026) Sigma- Aldrich | 1:1000 |
| Antibody | anti-Lamp-2 mouse monoclonal | (ab25631) Abcam | 1:500 |
| Antibody | anti-Vps26A rabbit polyclonal | (ab23892) Abcam | 1:2000 |
| Antibody | anti-RILP rabbit polyclonal | (ab128616) Abcam | 1:1000 |
| Antibody | anti-FLAG rabbit polyclonal | (F7425) Sigma-Aldrich | 1:1000 |
| Antibody | anti-EGFR rabbit monoclonal | (ab52894) Abcam | 1:1000 |
| Antibody | anti-LC3 rabbit monoclonal | (ab192890) Abcam | 1:2000 |
| Antibody | anti-CD63 mouse monoclonal | (556019) BD Biosciences | 1:500 |
| Antibody | anti-sortilin polyclonal rabbit | (ab16640) Abcam | 1 μg/ml |
| Antibody | anti-CI-MPR mouse monoclonal | (ab2733) Abcam | 1 μg/ml |

